# Colorful Lizards and the Conflict of Collection

**DOI:** 10.1101/2023.08.10.552819

**Authors:** Colin M. Goodman, Natalie M. Claunch, Zachary T. Steele, Diane J. Episcopio-Sturgeon, Christina M. Romagosa

**Affiliations:** Department of Wildlife Ecology and Conservation, University of Florida, Gainesville, Florida, 32611, USA; School of Natural Resources and Environment, University of Florida, Gainesville, Florida, 32611, USA; Department of Wildlife Ecology and Conservation, University of Florida, Gainesville, Florida, 32611, USA; Twitter: @heart2herp; School of Natural Resources and Environment, University of Florida, Gainesville, Florida, 32611, USA

**Keywords:** Attitudes, chameleon, human dimensions, indirect effects, non-native species, urban ecology

## Abstract

Invasive species threaten biodiversity and their management is economically burdensome. Research on the indirect effects of introduced species are often focused on indirect ecological effects, with little focus on the more difficult to capture but critically important societal impacts. Often understated are the social costs of invasive species such as conflicts between invasive species managers and public stakeholders. Chameleons are popular in the pet trade and have been introduced throughout Florida, and their presence often attracts private collectors. After locating a population of panther chameleons (*Furcifer pardalis*) within a suburban neighborhood in central Florida, we administered anonymous questionnaires to residents to explore how this introduction and the ensuing species collection has affected them. Respondents had knowledge of chameleon presence but most expressed low concern about chameleon presence. Respondents who had observed chameleons in the area expressed more concern for their safety given the activities of private collectors. Our study highlights the importance of recognizing the social impacts of species introductions in urban environments, particularly the attention these species can draw and the mixed perception of these species among stakeholders.

## Introduction

The global trade in wildlife products is flourishing, with more than 100 million CITES-listed (Convention on International Trade in Endangered Species of Wild Fauna and Flora) specimens traded annually (Harfoot et al. 2018). Legal trade in wildlife has more than quadrupled since the 1970s, and illegal sales account for an estimated $8–10 billion per year (Lawson and Vines 2014). In the United States, 224.9 million live animals are imported annually (Smith et al. 2017). Importation of wildlife is associated with the introduction and proliferation of non-native species (Lockwood et al. 2019). In few places is this more salient than Florida, where roughly 86% of known nonnative herpetofaunal introductions (n = 164) are attributed to the wildlife trade (Krysko et al. 2016). This has resulted in Florida having the largest number of established nonnative herpetofaunal species in the world (Krysko et al. 2016). The high demand for various herpetofaunal species encourages captive-breeding, and although captive-breeding is generally done under controlled conditions (Herrel and van der Meijden 2014), there is evidence of numerous intentional introductions of herpetofauna in Florida due to the hospitable climate (Krysko et al. 2003). One reason for intentional introduction is to generate a self-sustaining, free-ranging breeding population requiring minimal investment and upkeep (Krysko et al. 2003, Episcopio-Sturgeon and Pienaar 2019). Introduced populations may spread beyond the boundary of the initial introduction site, resulting in residents in these areas encountering the animals in their neighborhood or on their private property. Additionally, the rise of community science reporting platforms means that someone can publicly share information about an encounter with the species in question almost immediately. One such platform, the Early Detection and Distribution Mapping System (EDDMapS; Bargeron and Moorhead 2007), allows users to upload georeferenced observations of any non-native species they encounter.

Publicly available data from community science platforms make it increasingly difficult for the location of introduced populations to remain obscured long-term, and can be a cost-effective tool in non-native species management (Hester and Cacho 2017). However, these platforms rely on the voluntary participation of the public, which can be low. Additionally, cryptic species may still evade detection by willing residents. Further, hobbyists, breeders, and researchers alike monitor these platforms for hints of new populations (personal observation). When the location of a new population becomes public, an influx of people may seek to observe or collect animals from the population. In an urban or suburban environment, this is likely to result in interactions between the collectors and residents of the area. There is a growing body of evidence suggesting that public perception can impact the ability of biologists to effectively respond to and manage non-native species (Estévez et al. 2015). For example, negative perception of biologists or lack of trust can prevent eradication efforts if residents oppose biologists or are unwilling to allow access to private property to remove non-native species (Bertolino et al. 2021). Negative interactions between residents and private collectors may create additional challenges for research-based removal efforts, as otherwise cooperative residents may be unable to distinguish between individuals with private interests and those belonging to research organizations.

The willingness of residents to support or cooperate with management efforts is often contingent on the physical appearance of a species, and the direct and indirect effect of the species introduction (Bremner & Park 2007, Olszańska et al. 2016, Steele & Pienaar 2021). Direct effects include, but are not limited to, competition with native species, habitat destruction, disease transmission, and predation of native species (Gozlan et al. 2013). Species introductions often have indirect effects that are not as easy to decipher as the direct effects, and are therefore less studied. For example, the introduction of the brown tree snake (*Boiga irregularis*) devastated native bird species (direct) which resulted in dramatic loss of plant species that relied on the birds for seed dispersal (indirect; Rogers et al. 2017). Research on the indirect effects of introduced species are often focused on indirect ecological effects, with little focus on the more difficult to capture but critically important societal impacts (Reaser et al. 2007). Local residents may experience indirect societal effects associated with the introduction of a species, especially when that species is in high demand or is deemed to have aesthetic value (Bremner & Park 2007). Social impacts of introduced species include, but are not limited to, disruption of social activities, and induction of fear for safety (Bacher et al. 2017). Local residents may experience attachment to charismatic introduced species, making management efforts more difficult, or may experience increased disturbance due to a new presence of researchers or other parties interested in the introduced species. In Florida, individuals may experience indirect societal effects because of introductions of desirable non-native fish and herpetofauna species introduced for ranching throughout the state (Episcopio-Sturgeon & Pienaar 2019). This indirect effect often appears in the form of conflict between wildlife managers attempting to remove these populations and the public stakeholders who value these populations (Bertolino et al. 2021). In particular, the numerous pet reptile species introduced for ranching throughout the state of Florida may generate populations that local residents want to protect. Alternatively or concomitantly, as knowledge of the ranched population grows, interested individuals from surrounding areas may create local disturbances due to their collection efforts.

Given their low movement rates (Tolley et al. 2010), high value (Carpenter et al. 2004), and difficulty to maintain in captivity long-term (Stahl 1996), chameleons (members of the family Chamaeleonidae) are particularly suited for ranching. As a result, populations of numerous chameleon species have been intentionally seeded throughout the state of Florida (Gillette et al. 2010, Episcopio-Sturgeon and Pienaar 2019). Since these lizards have high value in the pet trade (Carpenter et al. 2004), populations are often kept secret by ranchers to prevent collection by other individuals. We became aware of an incipient panther chameleon (*Furcifer pardalis*) population limited to a few neighborhoods within central Florida from public posts made on social media by private collectors that had been in the area for the specific purpose of collecting chameleons. Chameleons are extremely difficult to observe in trees during daylight hours, but are easily distinguished from vegetation when spotlighted with flashlights at night. Thus, we conducted these surveys at night—when chameleons are inactive and less cryptic—by illuminating trees with spotlights (Tolley et al. 2010). This nighttime activity and atypical spotlight use (i.e., pointing upward at trees) may make the presence of collectors more obvious to residents in the area. As such, chameleons are an ideal species to study for the indirect societal effects of a species introduction.

We sought to determine how residents were affected both directly and indirectly by the presence of non-native species as well as the presence of the collectors, using an extant population of the non-native *F. pardalis* as a case study. To investigate sources of potential indirect conflict, we conducted an exploratory study by implementing a questionnaire to examine resident attitudes towards the nighttime activity (i.e., flashlight use) of collectors. Additionally, to distinguish potential impacts of the presence of non-native species from the presence of collectors, we evaluated resident knowledge and attitudes toward non-native reptile species (including chameleons) in this neighborhood. As such, we divided neighborhood residents based on responses: 1) Individuals who perceived the direct effects of the chameleon introduction; and 2) Individuals who perceived both the direct and indirect effects of chameleon introductions. We hypothesized that regions that had high concentrations of chameleons would be more likely to have higher numbers of chameleon sightings by residents, more likely to attract collectors leading to greater awareness of flashlights by residents, and higher levels of concern for safety from residents indirectly effected due to the presence of individuals with flashlights.

## Methods

### Study Site

The study region consists of a few small neighborhoods in Orange County, central Florida, encompassing an area of approximately 0.9 km^2^. To protect the privacy and identity of the neighborhoods, and to address the concerns expressed by residents during the questionnaire implementation, we have withheld specific locality information in this manuscript.

### Chameleon Detection Surveys

In this exploratory study, we performed two different types of surveys: one where authors sought and captured chameleons to determine presence (chameleon-detection surveys), and one where authors administered virtual questionnaires to residents (questionnaires). In late October 2019, we (and presumably others) found evidence suggesting the existence of the chameleon population on social media. We confirmed the presence of panther chameleons in the study area using systematic chameleon-detection surveys, documenting 26 total individuals (adults and juveniles) from 6 visits from 28 October–L10 November 2019. To confirm population presence prior to implementation of the questionnaire, we conducted an additional detection survey on 12 February 2020. We conducted chameleon-detection surveys at night with the use of personal spotlights. Surveys consisted of multiple observers walking bidirectionally along public walkways and using spotlights to illuminate present chameleons in trees. To minimize disturbance to the residents, we ceased all chameleon-detection surveys prior to midnight, carried university credentials, and only surveyed publicly accessible areas. Two specimens were collected and deposited in the Florida Museum of Natural History (Gainesville, Florida; UF 192232, UF 192233) as vouchers. All animals were collected with the approval of the University of Florida Institutional Animal Care and Use Committee, number 201910938. During these surveys, there were numerous instances where we observed private individuals surveying the area with the expressed intent of collecting chameleons.

### Questionnaire Design

We designed a questionnaire using Qualtrics (Qualtrics, Provo, Utah, USA), which varied in length depending upon the responses given (e.g., if participants reported having never seen chameleons, it did not ask for frequency of chameleon observations). All research was approved by the University of Florida Institutional Review Board, number 201903116, and all respondents provided their informed consent. We thoroughly pre-tested the questionnaire with Florida residents and both social science and ecological experts using both conventional and cognitive pre-testing. Pre-tests confirmed that all terminology was clear and that information presented in the questionnaire did not induce response bias (such as a respondent indicating multiple chameleon sightings because they believe this is socially desirable).

### Sample Population

Prior to questionnaire administration, we divided the study area into three regions: 1) core region; 2) external region; and 3) perimeter region (Fig. 3). These regions were determined by their proximity to the area chameleons were recorded during chameleon-detection surveys, with the core area being the region where we observed chameleons, the external region being the most proximal area (where chameleons were not detected by authors, but were by other collectors, so may exist in a much lower density), and the perimeter region being the farthest region from the known range where chameleons were not detected. The core region is separated from the external and perimeter regions by roadways, which served as natural boundaries for questionnaire boundary delineation. Each region received the same questionnaire, but each questionnaire was registered to its respective region to facilitate inter-region comparisons. Exploratory chameleon-detection surveys and questionnaire implementation resulted in no chameleon observations and no chameleon sightings reported from respondents in this perimeter region. Additionally, there were no reports of flashlights observed from respondents in the perimeter region. We thus excluded the perimeter region and focused our comparisons on the external and core regions (low vs high apparent density of chameleons, respectively).

This was a non-probability, intercept sample because we administered the questionnaire and distributed flyers with questionnaire links door-to-door in the area of focus. As such, the potential sample population depended upon who was present at the home and was willing to respond to our in-person request or noticed the flyers and had internet access. We visited and distributed flyers to 248 houses within the core and external regions, (Fig. 3; thus our potential maximum sample size from these regions was 248 because we only allowed one response per household. If a respondent stated they observed only chameleons, this respondent was deemed to only have experienced the direct effects of the chameleon introduction. If a respondent stated they observed chameleons and individuals with flashlights, this respondent was deemed to have experienced both the direct and indirect effects of the chameleon introduction (see Figure 1).

**Figure 1.**
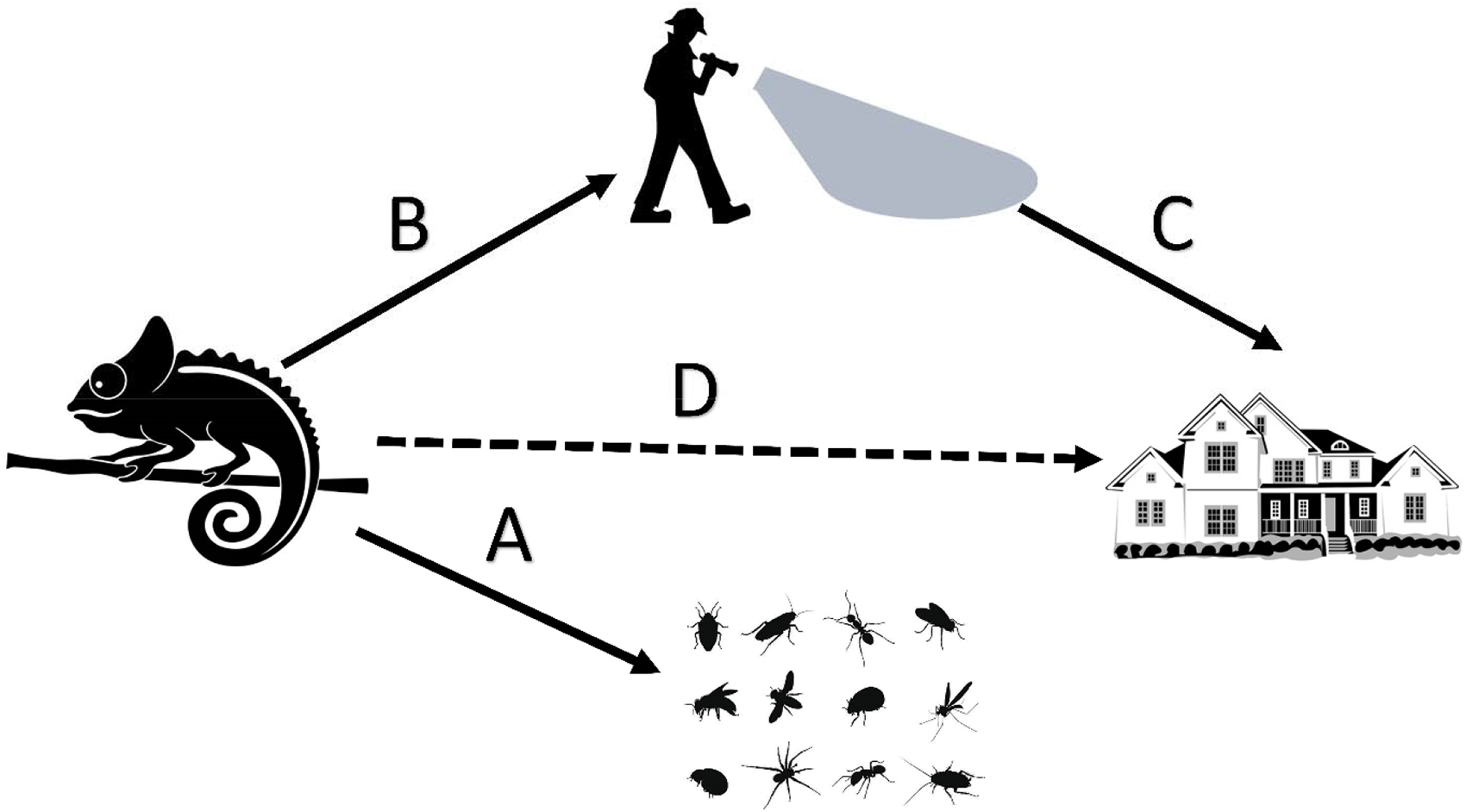
Visual showing the different effects of the chameleon introduction. Arrow A represents the direct effect of the introduction on native flora and fauna. Arrow B represents the direct effect of the introduction resulting in collectors flocking to the neighborhood to capture chameleons. Arrow C represents the direct effect of collectors on neighborhood residents. Finally, Arrow D represents the indirect effect of the introduction of chameleons resulting in neighborhood residents being concerned and disturbed by the presence of collectors. During sampling for the questionnaire, some respondents only reported their perception of the direct effects such as those highlighted via Arrow A. However, some respondents reported both the direct and indirect effects.

### Questionnaire Implementation & Analysis

We implemented the in-person questionnaire on 22 February 2020. We recruited participants by going door-to-door, starting at the core region and then moving outward to the external region. Participants answered the questionnaire on an iPad that we provided. If no one at a household responded, we left a flyer at the door with a description of the questionnaire and a corresponding QR-code and direct link to the questionnaire. Our data collection strategy changed in March due to the impacts of the COVID-19 pandemic, and we began solely distributed flyers throughout all regions through March 15. After this date, we ceased data collection due to concerns regarding COVID-19. We received the last recorded response from the flyers on 26 March 2020.

We first determined if respondents had observed panther chameleons in their area. Using unlabeled still images of reptiles, we prompted respondents to select reptiles they observed in their neighborhood. Along with images of male and female panther chameleons (to account for sexual dimorphism in color), we included an invasive reptile species common to the area (the brown anole – *Norops sagrei*) along with various visually iconic reptile species that are not native and have not been reported in the area (e.g., frilled-necked lizard – *Chlamydosaurus kingii*) to determine the integrity of responses.

If a respondent indicated observing a panther chameleon in their neighborhood, the questionnaire asked: 1) how long ago they first remembered seeing one; 2) how many times they had observed the species in the area; and 3) if they were aware that this species was non-native. For comparison, we asked these same questions to respondents who indicated they had observed the ubiquitous brown anole in their neighborhood. Questions for each species were accompanied by a photograph of the species to avoid confusion because anoles are sometimes colloquially referred to as “chameleons” due to their superficial resemblance to true chameleons (Carlton 1903). We then determined how concerned respondents were about the panther chameleons in their neighborhood using an ordinal scale (1-not at all to 5-extremely) on the following possibilities: 1) scratching/biting people; 2) transferring diseases to people; 3) harming [their] pet; 4) harming local wildlife; and 5) damaging [their] property. We again asked these same questions to respondents who indicated they had observed the anole in their neighborhood for comparison. We performed independent t-tests (interval responses), chi-square test (binary), or Mann-Whitney tests (ordinal responses) to test for differences between responses for the panther chameleon and brown anole.

To examine potential impacts of collectors with flashlights (indirect effects of chameleon introduction), the remainder of the questionnaire focused on the presence of individuals with flashlights in the respondents’ neighborhood. We first asked respondents: “Have [they] noticed flashlights shining around [their] neighborhood at night?”. If the respondent said “yes”, we documented when respondents first recall seeing flashlights and how frequently they have seen these flashlights at night. We then explored how seeing these flashlights made respondents feel in terms of being scared, frustrated, and curious, using the same five-point ordinal scale as above. We also determined if the people shining these flashlights impacted respondents’: 1) feelings of safety in [their] home; 2) feelings of safety in [their] neighborhood; 3) [their] ability to sleep; and 4) willingness to go outside at night. We used principal factor analysis to test for dimensionality of items that were intended to measure a specific composite variable (e.g., by summing individual items to generate scores that measured respondents’ described feelings from encountering individuals shining flashlights). We retained factors with an eigenvalue greater than one (using Kaiser-one criterion) and we report which items loaded onto retained factors (Santos 1999). We also ascertained the reliability and internal validity of items using Cronbach’s alpha at a threshold of 0.7. We examined if responses to these variables were significantly different depending on whether respondents who saw flashlights at night also had observed chameleons using Kruskal-Wallis tests. We examined if there was a correlation between observing chameleons and observing flashlights using Cramer’s V coefficient.

We asked respondents if they had ever interacted with the people shining flashlights. If the respondent stated “yes”, we asked if the individuals with flashlights indicated their purpose for being in the neighborhood. If the respondent stated that the person had indicated their purpose, we had the respondent describe the indicated purpose from the following options: 1) searching for wildlife; 2) collecting wildlife for sale; 3) collecting wildlife for research; and 4) unclear purpose / I don’t remember. This question allowed respondents to select multiple purposes. We also included an open-ended response if the purpose was not included in those options. We then asked respondents if they had reported the activities involving these shining flashlights to the police, their homeowner’s association (HOA), their neighbors, or ‘other’. This question also allowed selection of multiple answers and the option to provide an open-ended response to provide details about reporting these activities to another entity.

Respondents that spend more time outdoors may be more likely to encounter chameleons or chameleon collectors with flashlights, which may bias our sample. To evaluate this, we asked respondents how often they: 1) spend time gardening; 2) spend time outside with family/pets; 3) walk around their neighborhood/walk their pets; 4) exercise outdoors; and 5) observe wildlife outside (five-point ordinal scale, never to six–seven days per week). We used the Kruskal-Wallis test to determine if there was a significant difference in reported amount of outdoor activity depending whether the respondent reported observing a chameleon. All data were analyzed using Stata/SE 15.0 (StataCorp 2017) and R studio/R version 3.6.1 (RStudio Team 2020, R Core Team 2017).

To evaluate resident knowledge of wildlife reporting options, we asked respondents if they were aware of resources available to report the sightings of wildlife (EDDMapS, iNaturalist, HerpMapper, and IveGot1). The objective was to determine if respondents who had seen chameleons knew where they could report these sightings. We provided an open-ended textbox at the end of the questionnaire to allow participants to make additional commentary.

## Results

### Demographics

We obtained responses from 33 individuals in the study area, with the majority of responses coming from the core region (n = 22). We also obtained 11 responses from the external region. Overall, our response rate was 13% (n = 33), although our response rate for the core region was slightly higher (17%). The median age range of respondents was 45–54 and the majority of respondents reported living in the neighborhood for over ten years, with the median duration being 16.5 years.

### Awareness, Knowledge, and Concerns about Chameleons

There were no respondents in the external region that reported observing chameleons. Within the core region, 31.8% (n = 7) respondents reported observing chameleons at least once. Of the respondents who had previously seen chameleons, roughly half (n = 3) reported observing multiple chameleons for more than a year. There was a significant difference in knowledge of the non-native status of brown anoles and chameleons (ᵡ*^2^* = -1.97; *P* = 0.024); of the 30 respondents that reported seeing brown anoles, only 39% were aware of their non-native status. However, 86% of the respondents who reported seeing chameleons knew of their non-native status. Very few people reported concerns about the presence of either brown anoles or chameleons, and there were no significant differences in level of respondents’ concerns between the two species (Figure 2). Only two respondents were aware of any of the wildlife reporting platforms, and neither of these respondents indicated they observed chameleons in the area.

**Figure 2.**
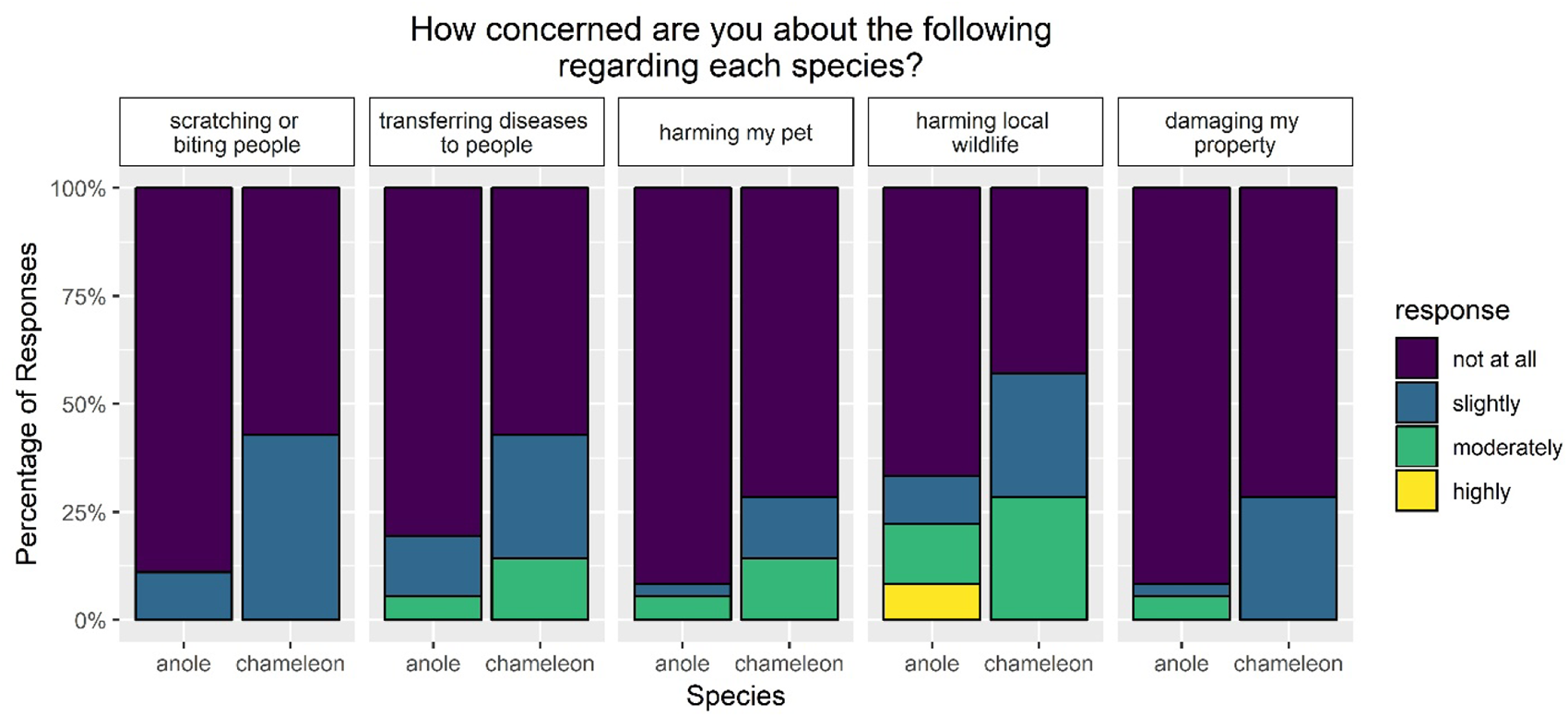
Differences in respondents’ perceived risk between brown anoles (n = 36) and chameleons (n = 7). Each paired panel represents a different cause of perceived risk. Each panel represents a different scenario, and each panel is split between anoles (left) and chameleons (right). Colors represent the level of concern expressed by respondents, and the height of each color represents the percentage of respondents that expressed that level of concern for each question.

**Figure 3.**
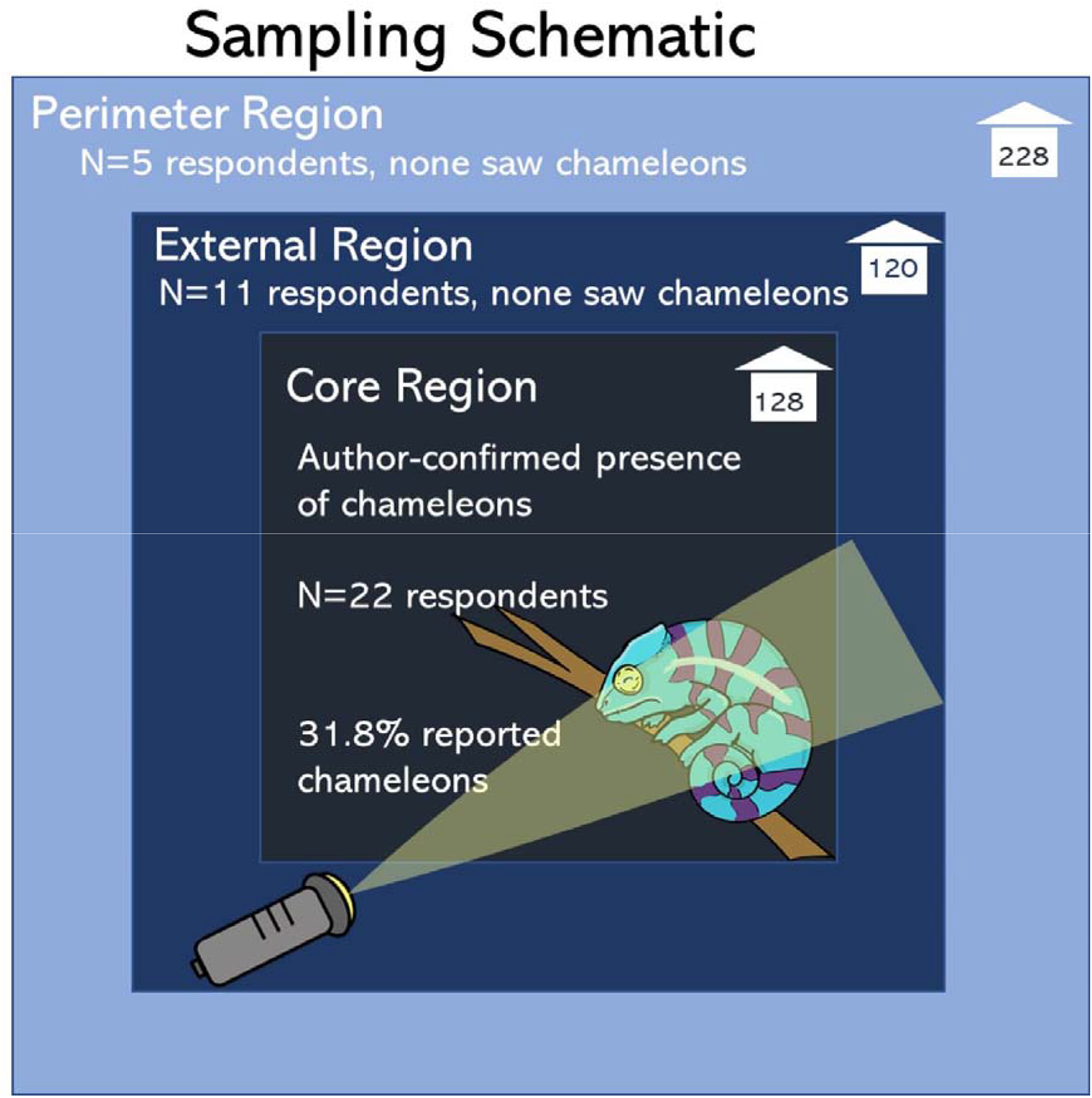
Sampling schematic illustrating the summary of responses and chameleon sightings reported from each region. Houses in the corner of each box represent the sampling effort in each region. Flashlights were reported from the external and core regions, while chameleons were only reported from the core region.

Using principal factor analysis, we generated a composite score for respondents’ reported time spent outside. This score was skewed left (mean = 13.56; range: 4–20), signifying that overall, respondents spent a large amount of time outside every week. There was a significant difference in the time spent outside between individuals who reported seeing chameleons and those who did not (χ^2^ = 5.19; df = 1; P = 0.022; Table 1).

**Table 1.** Kruskal-Wallis analysis of relationship with sightings of chameleons. Respondents were more likely to report sightings of chameleons if they spent more time outside every week. Respondents were more likely to report perceived feelings and impacts from seeing flashlights at night if they reported sightings of chameleons.

| Variables | $\chi^2$ | df | *P* |
| --- | --- | --- | --- |
| Time residing in neighborhood | 0.849 | 1 | 0.357 |
| Time spent outside | 5.191 | 1 | 0.022 |
| Emotional response to seeing flashlights<sup>a</sup> | 4.800 | 1 | 0.027 |
| Impact of seeing flashlights<sup>b</sup> | 4.640 | 1 | 0.031 |

<sup>a</sup>Denotes the composite variable derived from how scared, frustrated, and curious respondents reported feeling as a result of seeing flashlights.
<sup>b</sup>Denotes the composite variable derived from how seeing flashlights influenced respondents' perceived safety, ability to sleep, and willingness to go outside at night.

### Awareness, Knowledge, and Concerns about Collectors

Of the 33 respondents, 11 reported seeing flashlights shining at night in their neighborhood. Respondents who reported seeing flashlights shining at night were more likely to have also reported seeing chameleons (χ*^2^*= 12.46; φ = 0.558; *P* < 0.001), and approximately half of the respondents from the core region reported having seen both chameleons and flashlights shining at night. Only one respondent outside the core region reported seeing flashlights. There were no significant differences between the time when flashlights were first seen and the frequency with which the flashlights were observed. There was a significant difference in respondents’ emotional response to flashlights (χ*^2^* = 4.80; df = 1; P = 0.027; Table 1) and level of reported impacts from flashlights (χ*^2^* = 4.64; df = 1; P = 0.031; see discussion of both of these composite variables below) in their neighborhood if they reported seeing chameleons. Of the 11 respondents that reported seeing flashlights, more than half (n = 7) reported interacting with the individuals. One respondent reported having a confrontational interaction with individuals who repeatedly entered their yard for the purpose of collecting the panther chameleons. Another respondent reported approaching a group of collectors and calling the police after witnessing the collectors trespassing on a neighbor’s property (anonymous resident, personal communication). Among the respondents who reported approaching individuals with flashlights, a majority (n = 5) stated that these individuals indicated they were searching for wildlife and four respondents stated that these individuals indicated they were collecting wildlife for research purposes. Only one individual indicated they were collecting wildlife for sale. While only 5 of the 11 respondents who indicated seeing flashlights stated they reported these individuals, 80% of these respondents also indicated they observed chameleons. Four respondents indicated reporting individuals to a neighbor, and two reported contacting the police.

Using principal factor analysis, we created two composite variables. The first composite score (‘Emotional response to seeing flashlights’) combined respondents’ described feelings from observing flashlights shining and related to how scared, frustrated, and curious they felt as a result of this event (eigenvalue = 1.26; Cronbach’s alpha = 0.94). This first score was skewed right (mean = 8; range: 3–15), indicating that respondents did not have strong feelings about observing flashlights shining in their neighborhood. Respondents who observed chameleons and flashlights (n = 6) had an average emotional response score of a 10 while those who only observed flashlights average score was a 6 (n = 5). The second composite score (‘Impact of seeing flashlights’) combined respondents’ described impacts from observing flashlights shining and related to their perceived safety, ability to sleep, and willingness to go outside (eigenvalue = 3.19; Cronbach’s alpha = 0.71). This second score was also skewed right (mean = 7.55; range: 4–20), implying that respondents did not perceive many impacts from observing flashlights shining in their neighborhood. Respondents who observed chameleons and flashlights (n = 6) had an average impact score of a 10 while those who only observed flashlights average score was a 5 (n = 5).

### Limitations

The nature of this study is exploratory due to both a relatively small sample size and our planned efforts being extinguished by the impacts of COVID-19. Additionally, while residents did express concerns about seeing flashlights, the authors’ use of systematic chameleon detection surveys makes it impossible to determine whether all concerns arose due to the activities of private collectors, or whether some concerns arose due to the chameleon detection surveys of the authors. Regardless, this underscores the importance of public involvement to mitigate the impacts that management activities can have on residents.

## Discussion

This exploratory study represents the first attempt to evaluate potential indirect societal effects of introduced species. We sought to measure the awareness of suburban residents of a population of a highly sought-after nonnative species and the indirect societal effects associated with recent popularization of this population of animals (e.g., an increase in unfamiliar individuals in the area seeking out *F. pardalis*). Our results suggest that the focal population of *F. pardalis* is relatively localized. During our chameleon-detection surveys, we did not detect chameleons outside of the core region. Our surveys were not exhaustive (i.e., we did not survey private property within the regions except in instances when we were given explicit permission by the landowner). Thus, our lack of detections outside the core region may be attributed to a lower relative abundance of chameleons in these regions rather than true absences. For example, one non-resident collector we spoke with claimed to have captured chameleons well into the perimeter region of our study several years prior (anonymous collector, personal communication). However, we ultimately excluded the perimeter region from our analysis due to no respondents observing chameleons or flashlights, so no further analysis could be conducted. The questionnaire results supported the general findings of our detection-surveys because only respondents in the core region reported observing chameleons. Our results suggest that the *F. pardalis* population has occurred there for a minimum of two years, and conversations with residents outside of data collection suggest the population may be 10 years or older (anonymous respondent, personal communication). This was further corroborated by multiple private collectors, who stated they had been collecting chameleons from that area for over two years (anonymous collectors, personal communication). Despite the apparent age of invasion and prevalence of knowledge among respondents regarding the non-native status of *F. pardalis*, this population was previously not reported by residents, perhaps due to a lack of awareness of non-native species reporting platforms.

We found that very few respondents reported being concerned about the direct effects of the chameleon introduction, despite knowing their non-native status. There is evidence that individuals are highly influenced by invasion risks associated with their personal safety, livelihood, or the safety of a pet (Bacher et al. 2017, Steele & Pienaar 2021). Given their cryptic and arboreal nature, people may perceive chameleons more broadly as not posing a direct risk, even if less salient ecological risks exist. It is possible that respondents would not have reported chameleons to a community science platform even if familiar with one. We spoke with one resident who expressed an interest in having the chameleons protected at the state level (anonymous resident, personal communication), and even expressed a willingness to release chameleons found on public land onto their own property to preclude removal efforts. This sentiment may be exacerbated by the fact that unlawful, release of non-native species typically goes unpunished (Krysko et al. 2011). Lack of enforcement may remove any deterring impact a statute might have (Kahler et al. 2021) and may incentivize resident actions that will likely be at odds with the goals of wildlife managers, who are often focused on prevention of establishment, containment, or eradication of non-native species (Reaser et al. 2020).

While the chameleons themselves did not appear to directly impact residents, we documented indirect effects of chameleon presence via the actions of collectors surveying for them. We found that the presence of individuals with flashlights (i.e., potential chameleon collectors) appeared to indirectly impact and elicit stronger emotional responses in respondents who had seen both chameleons and flashlights. Residents who indicated they observed both chameleons and individuals with flashlights expressed concerns about safety in their homes, safety in their neighborhood, and feeling frustrated because of these shining lights at night. Related, respondents who had observed flashlights but had not seen chameleons were less likely to be affected by the flashlights. This may simply be because these respondents assumed the flashlights were from neighbors walking nearby, whereas respondents who had seen chameleons—and potentially knew about their value to collectors—were more likely to be concerned with the presence of strangers. Additionally, the majority of people with flashlights who responded to resident inquiry claimed to be collecting chameleons for research purposes, with only one respondent reporting a collector explicitly stating their intention to sell chameleons. Between posts on social media and in-person encounters during surveys, we observed at least five individuals who were collecting chameleons from the area with the specific purpose of selling them. A potential source of bias is that individuals with flashlights may have withheld their true intentions, avoided interacting with residents, or simply responded affirmatively when approached by residents asking if they were “with the university research group” to avoid potential conflict. However, it remains a possibility that some proportion of the interactions noted in the questionnaire data may have been with the authors from several months prior. Rather than invalidating our results, this highlights an area of concern for future research and invasive species management-namely how to minimize impacts to residents during urban wildlife research and how to distinguish research from commercial activity when there are conflicting interests and multiple parties involved. Furthermore, research and invasive species management may result in perceived feelings of fear or frustration from the public that negatively impact management efforts (Bacher et al. 2017). In situations in which local members of the public are experiencing social impacts of non-native introductions, the increased presence of unknown individuals seeking out the non-native species may result in negative attitudes toward the management of species (Bertolino et al. 2021). Local residents may prefer the presence of the non-native species in place of the disturbances they may associate with managing the non-native species (Bertolino et al. 2021, Steele – Accepted).

Mismatch between the goals of managers and other stakeholders can lead to problems with managing invasive species populations (Gozlan et al. 2013), often a result of the indirect societal effects of non-native introductions. While non-native species that have—or are perceived to have—a direct negative impact on people or property may garner increased support for direct management action (Olden and Tamayo 2014), the resulting disturbances of individuals looking for non-native species (whether an increased presence of wildlife managers, scientists, or non-native species collectors) may result in a social disturbance to local residents’, potentially disrupting social activities or inducing uncertainty and fear (Bacher et al. 2017). This social impact of non-native species may impact stakeholders’ attitudes toward non-native species management. This indirect effect of a species introduction may lead to public residents restricting access to their private property or sabotaging collection methods utilized by wildlife managers to remove a non-native species (Bertolino et al. 2021, Steele – Accepted). This emphasizes the importance of clearly communicating the direct effects of a species introduction to public stakeholders (Steele & Pienaar 2021). While it is unclear what impact *F. pardalis* is having on native species (in part due to the difficulty of locating individuals or populations for such studies), the Jackson’s chameleon (*Chamaeleo jacksonii*), which has invaded Hawaii, has demonstrated adverse impacts on native fauna (Kraus et al. 2012). Despite this precedent, residents may not be as supportive of management efforts if the direct effects of chameleons are not yet demonstrated, the direct effects do not personally impact residents, or if residents have a positive view of the species (Bacher et al. 2017, Beever et al. 2019, Steele & Pienaar 2021). Increasing public knowledge about non-native species may increase public support for invasive species management (Novoa et al. 2017, Steele & Pienaar 2021), but managers must recognize and tailor outreach towards a diversity of values and attitudes depending on the target species and the proximity of the species’ population to urban or suburban areas.

Despite best practices when collecting specimens (wearing university credentials, limiting survey hours, surveying only public areas, avoiding the pointing of flashlight beams towards residences), we likely created disturbances within the study area. This reflects the difficulties of data collection as wildlife continues to become less truly “wild” and more urbanized due to expanding human populations. The negative emotions and perceived impacts of the presence of individuals with flashlights should not be understated. Chameleons in this area appear to occupy both public and private property, and this may present a problem to managers. Private lands can serve as refugia for the target species, and failure to gain access to private lands can act as a hurdle to eradication efforts (Gardener et al. 2010, Witmer and Hall 2011). Determining whether a species should be managed should not only be determined by the risks presented, but also by evaluating the willingness of public to cooperate with the proposed management actions, especially if a large portion of the invasive range covers private property (Bertolino et al. 2021, Steele – Accepted). Encouraging cooperation may require elaborate and detailed campaigns, such as door-to-door approaches (Baldacchino et al. 2016). Evaluating the likelihood of stakeholder participation is critical given the high cost of even relatively small management operations. For example, a failed effort to eradicate a small population of Mexican gray squirrels (*Sciurus aureogaster*) in Florida was estimated to cost ∼$70,000 despite a small invasive range of less than three hectares and removing fewer than 100 individuals (Pernas and Clark 2011). While the level of conflict related to this specific situation appears low, negative interactions between the public and researchers, or collectors more broadly, can erode at public trust and increase the level of conflict (Zimmerman et al. 2020). Without this trust, the public may be less likely to support, and therefore cooperate with, management efforts (Estévez et al. 2015).

It is imperative that researchers actively engage with the stakeholders that live in proximity to research sites. Cerra (2017) suggests community involvement as a way of fostering biodiversity stewardship among private landowners. This strategy is likely to require organizing public events or canvassing efforts to tell residents about the research being conducted in their area. Dyson et al. (2019) recommends engaging with and having contact information of a superior at the ready if approached by residents. When stakeholders are not included in planning for an invasive species management action or feel strongly opposed to the decision to a manage a species, there is the chance for serious conflict (Witmer and Hall 2010, Engeman et al. 2018). An effort to remove the Gambian giant pouched rat (*Cricetomys gambianus*) from Florida was partially disrupted when local residents directly sabotaged traps (Witmer and Hall 2010). Florida residents may have also reintroduced feral swine (*Sus scrofa*) to the island of Cayo Costa in 2008 in direct opposition to management efforts just weeks after an apparent successful eradication (Engeman et al. 2018). While these examples encapsulate a higher level of conflict than the current panther chameleon situation (Zimmerman et al. 2020), they highlight the importance of community involvement, and these recommendations are congruent with our own findings. Many of these residents we spoke with during chameleon-detection surveys expressed an interest in the research and a willingness to help. This suggests that management of the panther chameleon population may still be feasible at the present level of conflict (Zimmerman et al. 2020), and indicates a mismatch between the values professed by residents and the requisite knowledge to adhere to those values, and likely stems from respondents’ lack of knowledge of available reporting platforms.

Survey evidence from the public as well as community science experts suggests that awareness of community science initiatives more broadly is quite low (Lewandowski et al. 2017). Similarly, outreach efforts at informal education centers such as zoos and aquariums pertaining to community science initiatives for reporting invasive species sightings appears to be minimal (Steele and Pienaar 2023). Despite the observed knowledge gap, there is evidence suggesting that outreach—in the form of public workshops—can increase public participation in these community science platforms (Wallace et al. 2019). Training people in reporting via community science platforms is also associated with increased knowledge and awareness of invasive species (Jordan et al. 2011, Crall et al. 2013, but see Brossard et al. 2005), as well as increased participation in conservation-related activities (Lewandowski and Oberhauser 2017). Thus, the dissemination of information on the existence and use of these platforms may also help bridge gaps between the values of residents and values of biologists and managers. If these hurdles can be overcome, nonnative species reporting platforms may be especially important for species that are not perceived to be hazardous to humans, and thus attract less media attention than species that are—often unjustifiably so—portrayed as and perceived to be hazardous to humans (Reed and Snow 2014, Moloney and Unnithan 2019).

## Conclusion

We recorded greater perceived indirect effects from the individuals using flashlights (chameleon collectors) in their neighborhood than directly from the non-native species of focus. A cognizant and willing public is a vital component to managing non-native species, particularly in urban settings. We found evidence indicating both a lack of knowledge among respondents of available community science platforms, as well as regarding the non-native status of the widespread brown anole. Lastly, we found evidence that the presence of collectors—including those that explicitly stated research-based intentions—negatively affected some respondents. Taken together, our results show a need for careful consideration when developing non-native species management programs in urban or suburban areas. Although the current panther chameleon population appears isolated, failure to engage with residents and appeal to their values will likely result in an inability to manage this population properly. Overall, active management of non-native species can lead to conflicts that may be heightened in urban areas. These social conflicts underscore the importance of increasing public education and engagement in the planning and execution of non-native species management.

## Acknowledgements

We would like to thank C.D. Raymond, B.L. Phillips, P. Miller, M. Sandfoss, K. Hengstebeck, D. Juárez-Sánchez, M. Vilchez, S. Nielsen, S. Tillis, D. Catizone for their contributions to this research.

